# Studies on the interaction of three lytic bacteriophages with a wide collection of *Escherichia coli* strains implicated in swine enteric colibacillosis

**DOI:** 10.1101/2022.07.04.498734

**Authors:** Alice Ferreira, Carina Almeida, Daniela Silva, Isidro García-Meniño, Azucena Mora, Hugo Oliveira, Ana Oliveira

**Author notes:** Alice Maria Fernandes Ferreira Carina Ferreira Almeida Daniela Sá Pereira e Silva Isidro García-Meniño Azucena Mora Gutiérrez Hugo Alexandre Mendes Oliveira Ana Cristina Afonso Oliveira. Corresponding authorE-mail (AO).

## Abstract

The misuse of antibiotics in the swine industry and their on-going restriction requires alternatives to control enterotoxigenic and shiga toxin-producing *Escherichia coli* (ETEC and STEC, respectively). This study evaluates the potential of three coliphages, vB_EcoM_FJ1, vB_EcoM_FN and vB_EcoM_SP1 against 104 ETEC, STEC and ETEC/STEC strains isolated from pig colibacillosis in Portuguese (2018-2020) and Spanish farms (2006-2016), encompassing 71.2% *mcr*-positive strains (33.7% with *mcr-1*, 1.9% *mcr-2*, 35.6% *mcr-4* and 2.9% *mcr-5*) and 18.3% positive strains for TEM (1%), SHV (6.7%), and CTX-M (11.5%) extended-spectrum beta-lactamase-encoding genes. In general, all bacteriophages presented a narrow lytic spectrum (up to 2.9%) against the 104 ETEC, STEC and ETEC/STEC. Bacteriophages shared >80% overall nucleotide identity with *E. coli* phage T4 (*Tevenvirinae* subfamily), but a particular look at the distal part of the long tail fiber (gp38) revealed no homology. All bacteriophages recognize lipopolysaccharides as receptors, and additionally, FN binds to an outer membrane protein A. Bacteriophage-insensitive mutants of vB_EcoM_FJ1 (90%) and vB_EcoM_FN (100%) were shown to be more susceptible to pig serum inactivation comparatively to the parental strain and furthermore, their adhesion capacity to porcine intestinal cells was diminished by, approximately, 90%. Contrariwise, vB_EcoM_SP1 insensitive variants did not display phenotypic differences comparing to the wild-type strain. This study demonstrates that besides being T4-like, these bacteriophages revealed a narrow lytic spectrum against diarrhoeagenic *E. coli* strains and that the acquisition of novel bacteriophage-encoded adhesins (gp38) seems to be determinant for such results.

## Introduction

Intestinal *Escherichia coli* associated infections are recurrent in pig farms worldwide and often originate from environmental contamination (i.e., wastewater and animal faeces). Enterotoxigenic *Escherichia coli* (ETEC) is the most prevalent pathotype involved in enteric colibacillosis outbreaks of neonatal and post-weaning diarrhoea, causing high rates of morbidity and mortality and requiring expensive control measures [1]. Animals with a fragile immune system, particularly in the neonatal and PW periods are more susceptible to the disease [2]. ETEC strains exhibit different colonization fimbriae that enable bacterial adhesion by recognition of specific receptors present in enterocytes. F4 and F18 are the most prevalent in the PW phase while F5, F6 and F41, whose recognition sites in enterocytes decrease with the age of the pig, are less frequent [1]. ETEC also produce heat stable (ST) and labile (LT) enterotoxins, responsible for overproduction of electrolytes and fluids, and reduction of water adsorption, causing acute diarrhoea, dehydration, slow growth and even death in pigs. Two types of ST enterotoxins, STa and STb were so far reported [3]. Shiga toxin–producing *E. coli* (STEC) also implicated in enteric colibacillosis, carries the shiga toxin type 2e (Stx2e) and adheres to enterocytes mainly through F18 fimbriae [1]. Hybrid strains (ETEC/STEC) are also observed [4].

The massive use in swine of last resort antibiotics used to treat humans (e.g extended-spectrum cephalosporins and colistin) has led to the presence of residues of extended-spectrum beta-lactamase-producing and multidrug resistant (MDR) *E. coli* in farms [5]. The European Medicine Agency has therefore restricted the use of antibiotics in farms, to mitigate the potential cross-contamination risks of resistant strains along the food chain [6]. Consequently, the reduction of available antibacterial options turns urgent the development of effective and sustainable alternatives. Some preventive measures are used to limit the impact of PW diarrhoea. The effectiveness of hygienic measures and strict biosecurity rules, such as vaccination of sows, use of prebiotics and probiotics, or genetic breeding for ETEC-resistant herds, although important, fail to avoid the use of antibiotics [1]. Bacteriophages (phages) are specific and obligatory bacterial parasites with a genome confined in a protein capsid. They vary on lifestyle (virulent and lysogenic), genome type (single and double stranded DNA or RNA) and morphology (mostly, tailed viruses are from *Myoviridae, Siphoviridae* or *Podoviridae* families). Their self-replicating, self-dosing capability and innocuity nature towards animal cells as well as their high specificity towards the target bacterium (not affecting the commensal microbiota) are valuable traits encouraging its use [7]. Virulent phages that replicate within the bacterial host, releasing their progeny after cell lysis, have been of particular interest to use against bacterial pathogens, including ETEC and STEC. Despite the proof of concept of phage efficacy in veterinary medicine is being reported [8], studies in pigs are still few. Yet, three studies have reported successful results in using phages both prophylactic and therapeutically to fight against few ETEC serotypes causing infections (O149:H10:F4 [9,10] and F4 carrying strains [11]).

This study brings new data and promote discussion about current bottlenecks on the successful use of phages to control swine colibacillosis. Here, we characterize three coliphages against a collection of Portuguese and Spanish *E. coli* strains implicated in swine enteric colibacillosis.

## Materials and methods

### Bacterial strains and culture conditions

In this study, 156 *E. coli* strains were isolated from pig farms in the North-Central region of Portugal pig farms between 2018 and 2020. The strains were collected from fecal samples and rectal swabs of pigs with diarrhoea aged between two days and one month old, or from intestinal contents of dead infected animals. The Portuguese collection was tested to detect the presence of ETEC and/or STEC pathotypes, as described below. Besides, a collection of 68 Spanish strains fully typed comprising 57 ETEC, five STEC and six ETEC/STEC representative of different and prevalent seropathotypes implicated in enteric colibacillosis, was included here [5]. As control, 36 Avian pathogenic *E. coli* (APEC) previously isolated from organs of infected chickens recovered in Portuguese avian farms, were used to comparatively assess the efficacy of the phages among diarrhoeagenic versus extraintestinal pathogenic *E. coli* (DEC/ExPEC, respectively). All bacterial strains were cultivated in MacConkey agar (50 g.L^−1^, Biokar Diagnostics) for isolation, grown in Lysogeny Broth (LB, NZYTech) agar (12 g.L^−1^, VWR) at 37 °C for the subsequent studies, and stored at −80 °C.

### IPEC-1 cells maintenance

For tissue cultures, the neonatal intestinal porcine cell line IPEC-1 (CVCL_2245) was used. Cells were maintained in Dulbecco’s Modified Eagle’s Medium (DMEM, Biochrom) and Ham’s F-12 (Biochrom) (1:1) supplemented with 10% fetal bovine serum (FBS, Biochrom) and 1x ZellShield (Biochrom) at 37 °C in a humidified atmosphere at 5% CO_2_ (HERAcell 150). IPEC-1 cells were subcultured every three days at 80% confluence in T-flasks (Starstedt) in 10 mL complete cell culture medium. Cells used in this study were subcultured from passage 12 to 17.

### Genotypic characterization of DEC

The *E. coli* pathotypes and virulence factors associated with enteric colibacillosis were investigated among the Portuguese swine strains by PCR of specific genes encoding for toxins (STa, STb, LT, Stx2e) and adhesins (F4, F5, F6, F18 and F41). Primer pairs and PCR conditions were previously reported by García-Meniño and colleagues [5] (S1 Table).

Those DEC strains conforming ETEC and/or STEC pathotypes were further analysed for the presence of colistin resistance associated to *mcr* genes (*mcr-1, 2, 3, 4* and *5*), using reported PCR conditions [5] (S2 Table). Then, the *mcr*+ strains were also screened for the detection of TEM, SHV, and CTX-M beta-lactamase-encoding genes [12] (S2 Table).

### Phylogroups, Sequence types (STs) and Clonotypes

The main phylogenetic groups of *E. coli* (A, B1, B2, C, D, E, and F) were determined for the *mcr*+ strains using the quadruplex PCR method described by Clermont et al. (2013) [13], based on the presence/absence of the four genetic targets *arpA, chuA, yjaA*, and *TspE4*.*C2* (S3 Table). The STs of the strains were assigned by multilocus sequence typing (MLST) following the Achtman seven-locus scheme [14] (S3 Table), and the allelic profile for each isolate was retrieved through the Enterobase website (https://enterobase.warwick.ac.uk/species/ecoli/allele_st_search). The clonotyping was based on the internal 469-nucleotide (nt) and 489-nt sequence of the *fumC* and *fimH* genes, respectively. Allele assignments for *fimH* were determined using the fimTyper database available at the Center for Genomic Epidemiology website (http://www.genomicepidemiology.org/). The combination of *fumC* (allele obtained from MLST) and *fimH* allele designations was used as the CH “type” [15] (S3 Table).

### O Typing

The most prevalent serogroups implicated in enteric colibacillosis of swine were investigated for the *mcr*+ strains by microagglutination, following the method described by Guinée et al. (1981) [16] and using the specific O45, O101, O108, O138, O139, O141, O149 and O157 antisera at the Laboratorio de Referencia de *E. coli* (LREC-USC). Strains that did not react with any of those O antisera were classified as non-assigned (NA) serogroup.

### Haemolysis type

The haemolytic capacity of ETEC and STEC strains was evaluated by observing the colony phenotype after cultivation in Columbia blood agar (BioMérieux) and incubated at 37 °C, overnight (O/N). The presence and type of lysis halos around the bacterial colonies identified alpha (α) (green discoloration around the colonies), beta (β) (clear zone or transparency in the surrounding medium) or gamma (ɣ) (absence of reaction, non-haemolytic) haemolysis.

### Antibiotic susceptibility

Antibiotic susceptibility was determined by microdilution assays and diffusion disks. Microdilution assays were performed to assess the phenotypic resistance to colistin. Briefly, O/N cultures were 100-fold diluted in fresh LB and gown at 37 °C, 120 rpm until mid-log phase. Bacterial suspension of an OD_600nm_ = 0.1 (1×10^8^ CFU.mL^−1^) were 100-fold diluted in colistin solution (final concentration 2 mg.L^−1^) in a 96-well polystyrene microplate (SPL Life Sciences). Next, the turbidity (OD_600nm_) was measured in a spectrophotometer (Heales, MB-580) after a 22 h incubation period at 37 °C. A bacterial suspension without colistin was used as a control. A *mcr*^−^ strain was used as positive control. The experiments were conducted in triplicate. Results were interpreted following the 2022 EUCAST breaking point (http://www.eucast.org).

Additionally, the strains were subjected to disc diffusion tests (Bio-Rad) containing gentamicin (10 µg), cefoxitin (30 µg), imipenem (10 µg), aztreonam (30 µg), amoxicillin + clavulanic acid (20 µg + 10 µg), ampicillin (10 µg), ceftiofur (30 µg), cefepime (30 µg), doxycycline (30 µg), minomycin (30 µg), colistin (10 µg), tigecycline (15 µg), marbofloxacin (5 µg), nalidixic acid (30 µg), ciprofloxacin (5 µg), enrofloxacin (5 µg), trimethoprim-sulfamethoxazole (75 µg) and fosfomycin (200 µg). An isolate was considered either susceptible, intermediate susceptible or resistant following the manufacturer guidelines based on CLSI breakpoints (M100 30th Edition, 2020).

When resistant to at least one agent in three or more antimicrobial categories, strains were classified as multidrug-resistant as proposed by Magiorakos et al. (2012) [17].

### Phage propagation

A panel containing three previously isolated phages were used. Phages vB_EcoM_FJ1 (FJ1) and vB_EcoM_FN (FN) were isolated from chicken litter (unpublished data). Phage vB_EcoM_SP1 (SP1) was isolated from pig sewage and previously reported [18].

For phage propagation the host cells, O/N grown host cultures - HFJ1, HFN and SP16, respectively for FJ1, FN and SP1 - were incorporated into LB soft agar overlays plates, before spreading phage suspensions (∼1×10^7^ PFU.mL^−1^) with a sterile strip of paper to produce confluent plates. Plates were then O/N incubated at 37°C, and after that, 3 mL SM buffer were added. Plates were again incubated (4 °C for 16 h), and then the liquid phase was recovered, centrifuged, treated with chloroform (10% v/v), filtered (0.2 µm) and stored at 4 °C. Phage concentration was assessed by plaque counting (PFU.mL^−1^) after serial diluting the phage stock in SM Buffer, plating and incubation.

### Electron microscopy analysis

Phages FJ1 and FN were visualized by Transmission electron microscopy (TEM). Phage particles (>1×10^8^ PFU.mL^−1^) were collected by centrifugation (1 h, 25,000 × *g*, 4 °C) and washed twice with water. Next, phages were deposited on copper grids with a carbon coated Formvar film grid and stained with 2% uranyl acetate (pH 4.0). The visualization was performed on a Jeol JEM 1400 (Tokyo, Japan).

### DNA isolation, genome sequencing and annotation

Genomic DNA of phages FJ1 and FN was isolated using the phenol-chloroform-isoamyl alcohol method as previously described [19]. The DNA sample was used for whole genome library construction using TruSeq® Nano DNA Library Prep Kit. DNA fragments were sequenced in Illumina MiSeq, using 300bp paired-end sequencing reads. After removing low quality bases, reads were *de novo* assembled using Geneious Prime. The assembled genomes were scan through MyRAST for open reading frames [20] and tRNAscan-SE for tRNAs [21]. Protein functions were search using BLASTP against NCBI nonredundant protein database and using HHpred against Protein Data Bank database, in all using a E-value 1×10^− 5^ threshold. Comparative genomic analysis was performed with BLASTN. Phages FJ1 and FN sequenced genomes were deposited in NCBI database under the accession numbers MZ170040.1 and MZ170041.1, respectively.

### Lytic spectra determination and efficiency of plaquing

The host range was evaluated against a wide panel of 104 DEC strains from pigs: 31 ETEC and five STEC isolated within the scope of the present work, and 68 ETEC, STEC and ETEC/STEC previously isolated in Spain [5]. Moreover, to assess phage activity against other *E. coli* pathogenic strains, 36 APEC strains, previously isolated from chickens were also included. Two parameters were then analysed. First, strains were subjected to phage spot test to assess the host recognition rate: 10 µL of each phage (1×10^8^ PFU.mL^−1^) were dropped onto each bacterial lawn (prepared as previously described) and checked for clear zones after incubation. Then, the range of plaque formation was evaluated in sensitive strains, by measuring phage efficiency of plaquing (EOP): serial dilutions of phage suspensions (starting from 1×10^8^ PFU.mL^−1^) were spotted on bacterial lawns. The relative EOP was calculated by dividing the titre (PFU.mL^− 1^) of each susceptible strain by the titre of the relevant propagating host, and scored as 0 (no lysis), 1 (≤50%), 2 (>50% - 100%), 3 (>100%) and lysis from without (LFW) if no single plaques were observed.

### One-step growth curve

One-step growth curves were performed for all phages (FJ1, FN and SP1). Shortly, O/N-grown cultures were 100-fold diluted in 20 mL of fresh LB and incubated until an OD_600nm_ of 0.3 was reached. Resultant cultures were then centrifuged (7,000 ×*g*, 5 min, 4 °C), resuspended in 5 mL fresh LB medium, and mixed with 5 mL of phage suspension to reach a multiplicity of infection of 0.01 (for FN) or 0.001 (for FJ and SP1). A subsequent incubation (37 °C for 10 min) allowed phage adsorption to bacterial cells and then a centrifugation (7,000 ×*g*, 5 min, 4 °C) produced a pellet that was resuspended in 10 mL of fresh LB medium. To analyse one-step growth curves, samples were taken every 5 or 10 min and plated immediately over a period of 35 min, 40 min or 50 min for FJ1, FN and SP1, respectively.

### Identification of phage receptors

The type of phage receptors (carbohydrate or protein-based) on bacterial surface was identified following the protocol proposed by Kiljunen et al. (2011) [22]. Phage host cultures were treated with 1) sodium acetate (50 mM, pH 5.2) (control), 2) sodium acetate containing 100 mM periodate (IO_4_^−^) at room temperature for 2 h (to inactivate carbohydrates) or 3) proteinase K (0.2 mg.mL^−1^) at 37 °C for 3 h (to inactive outer membrane proteins). Afterwards, the phage was incubated with the treated host cells during 5 or 10 minutes at 37 °C, and the adsorption measured by plaque counting (PFU.mL^−1^) after serial diluting in SM Buffer. The phage adsorption rate (%) was obtained by subtracting the concentration of non-adsorbed phage divided by the total phage titre. Each assay was performed at least 3 times.

Complementary studies were performed with phages FJ1, FN and SP1 to identify specific receptor-encoding genes, using the Keio Collection composed of *E. coli* K-12 mutants carrying single-gene deletions [23], performing drop tests.

### Bacteriophage-insensitive mutants’ survival in pig serum

The vulnerability of bacteriophage-insensitive mutants (BIMs) generated by phages to the pig complement system was assessed using porcine serum. For inducing the formation of BIMs, mid-log phase grown cultures of host strain EC43 were challenged with FJ1, FN and SP1, and incubated for 24 h. After incubation, the cultures were plated in LB agar and incubated O/N. Afterwards, 10 colonies obtained from each culture were streaked at least three times into new plates to guaranty purity. The confirmation of BIMs production was performed by EOP. Whenever 24 h of incubation were not enough to obtain insensitive mutants, the procedure was extended to 72 h.

Next, EC43 wild-type (WT) and respective BIMs mid-log phase cultures (OD_600nm_ = 0.3) were diluted to obtain a 5×10^5^ CFU.mL^−1^ and mixed with porcine serum (3:1 (v/v)). The mixture was incubated for 1 h at 37 °C, followed by quantification of bacterial cells. Heat-inactivated serum (56 °C for 30 min) was used as negative control.

### Phage-induced mutants’ adhesion to epithelial cells

The virulence of five BIMs of each phage (FJ1 - 1.1, 1.4, 1.6, 1.9 and 1.10, FN - 2.1, 2.2, 2.3, 2.7 and 2.9 and SP1 - 3.1, 3.5, 3.6, 3.7 and 3.9) in swine intestinal cells was assessed by comparing the adhesion capacity (CFU.cm^2^) caused by the BIM and by the originating strain. Briefly, IPEC-1 cells were seeded in 96-well plates and let growth for 24 h to confluence confirmed by microscopy. Afterwards, cells were washed once with 10 mM PBS and exposed to bacterial suspensions (MOI=100) resuspended in DMEM/Ham’s F-12 supplemented with 10% FBS and incubated during 2 h at 37 °C, 5% CO_2_ to allow adhesion. After incubation, the culture medium was removed, the cells were carefully washed twice with 10 mM PBS, 30 µL of trypsin/EDTA (Biochrom) was added to each well and plates were re-incubated at 37 °C, 5% CO2, for 15 min. The effect of trypsin was quenched by adding 70 µL of assay medium. CFU were quantified by 10-fold serial dilutions in 0.9% (w/v).

### Statistical analysis

The statistical analysis of the results was performed using GraphPad Prism 6. Results were compared using t-test or One-way ANOVA using Bonferroni test. All tests were performed with a confidence level of 95%. Differences were considered statistically different if *p*-value≤0.05.

## Results

### Virulence factors, serogroups, mcr-types and beta-lactamase encoding genes

A total of 156 *E. coli* strains were isolated from faecal samples or rectal swabs during diarrhoea outbreaks in Portuguese pig farms between 2018 and 2020. While most strains (76.9%) tested negative for all the virulent-related traits analysed by PCR, 36 (23.1%) of the strains could be encompassed in two pathotypes (ETEC and STEC). Of the 36 pathogenic strains, 86.1% carried enterotoxin genes (ETEC), of which 90.3%, 48.4% and 29.0% of the ETEC strains showed carriage of STb, STa or LT genes, respectively. The remaining 13.9% strains carried the shiga-like toxin gene *stx2e* (STEC) (Table 1). Regarding the intestinal colonization factors, the most prevalent fimbriae among ETEC was F18 (41.9%) followed by F4 (16.1%). Most ETEC strains carried both fimbriae and toxin-encoding genes (58.1%). Fimbriae F18 was also present in 80% of the Shiga-like toxin-bearing strains. Fimbriae F5, F6 and F41 were not detected within the collection. Additionally, 54.8% and 80% of ETEC and STEC strains respectively, displayed β haemolytic activity.

**Table 1.** Virulence factors (fimbriae, toxins) attributes of the 36 Portuguese DEC isolates and haemolysis type.

| Pathotype | Fimbriae<br>No. isolates (%) |  | Toxins<br>No. isolates (%) |  |  |  | Haemolysis type<br>No. isolates (%) |  |
| --- | --- | --- | --- | --- | --- | --- | --- | --- |
| | F4 | F18 | STa | STb | LT | Stx2e | $\beta$ | $\gamma$ |
| ETEC<br>(n=31) | 5 (16.1) | 13 (41.9) | 15 (48.4) | 28 (90.3) | 9 (29.0) | - | 17 (54.8) | 14 (45.2) |
| STEC<br>(n=5) | - | 4 (80) | - | - | - | 5 (100) | 4 (80) | 1 (20) |

The screening of plasmid-mediated colistin resistance genes (*mcr-1* to *5*) on the 36 ETEC and STEC identified 36.1%, 5.6% and 25% strains with *mcr-1, mcr-2* and *mcr-4*, respectively. Among the *mcr*+ strains, three different serogroups were identified. Predominantly, ETEC strains belonged to serogroups O108 (31.6%) and O157 (15.8%) while all STEC strains belonged to O139 (S4 Table).

The screening of the *bla*_*CTX-M*_, *bla*_*SHV*_, and *bla*_*TEM*_ genes within the 24 *mcr*+ strains bearing genes indicated that all but one strain (*bla*_*CTX-M*_ carrying) displayed *bla*_*TEM*_ in their genomes. One strain encompassed both *bla*_*TEM*_ and *bla*_*CTX-M*_ genes. No *bla*_*SHV*_ gene was identified (S4 Table).

### Phylogroups, STs, clonotypes

By PCR, the 24 *mcr*+ ETEC and STEC strains were assigned to four distinct phylogroups: A (15 strains), B1 (three strains), E (five strains) and F (one strain). MLST determined six different STs, but 15 strains of 24 belonged to CC10 (S4 Table). Among the seven phylogroup-ST-clonotype (CH) combinations determined within the 24 strains, three of them accounted for 79% of them: A-ST10 (CH11-24), A-ST5786 (CH11-24) and D-ST1 (CH2-54). Interestingly, all 9 *mcr-4* strains belonged to the clonal group A-ST10 (CH11-24), mostly exhibiting serogroup O108. Besides, the five STEC strains showed the clonal group D-ST1 (CH2-54), serogroup O139 and carried *mcr-1*.

### Antimicrobial resistances

The inhibition assay confirmed that the *mcr* carrying strains were resistant to colistin at 2 mg.L^−1^ (EUCAST breakpoint). Additionally, the antibiotic resistance profile (Fig 1) indicated high resistance rates to ampicillin (100.0%), trimethoprim-sulfamethoxazole (83.3%), doxycycline (75%), gentamicin (70.8%), nalidixic acid and ciprofloxacin (62.5%) and enrofloxacin (50.0%). Most strains (75% and 50%) displayed an intermediate susceptibility to colistin and amoxicillin + clavulanic acid, respectively. The active ingredients with higher effectiveness were fosfomycin (100.0%), cefoxitin (95.8%), imipenem (91.7%), cefepime and tigecycline (79.2%), aztreonam (75%), marbofloxacin (62.5%) and ceftiofur (54.2%) Also, based on the antibiotic resistance pattern, all 24 strains were considered MDR.

**Fig 1.**
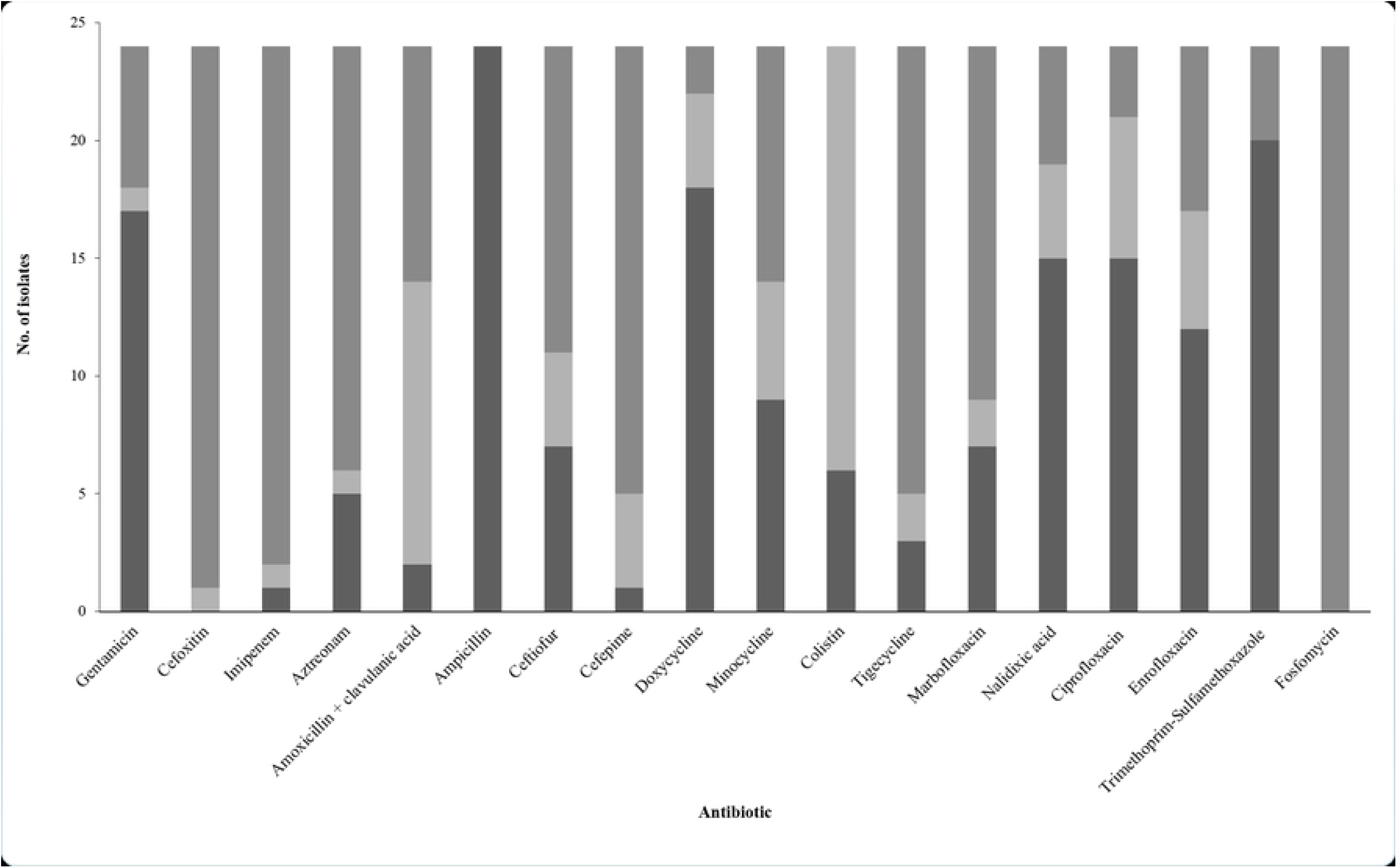
Antimicrobial susceptibility of the 24 *mcr*+ DEC Portuguese strains. The strains were assessed for their susceptibility towards a wide range of antibiotics used in the swine industry. Results were interpreted according to the CLSI, 2020. Grey colour stands for susceptible; light grey colour means intermediate susceptibility; dark grey colour stands for resistance.

### Bacteriophage isolation, host recognition and plaque formation efficiency

The lytic spectra and EOP of the three phages (FJ1, FN and SP1) were firstly tested against 88 ETEC, 10 STEC, six ETEC/STEC (S5a Table). Overall, all phages demonstrated a narrow lytic spectrum: FJ1, FN and SP1 were able to lyse and propagate, respectively, in 1.0%, 2.9% and 1.0% of the 104 ETEC, STEC and ETEC/STEC (from which only FJ1 - 100% - score an EOP greater than 50%). Additionally, phages recognized and lysed without propagation (LFW) 4.8%, 19.2% and 9.6% of the same strains.

The EOP was further performed in 36 APEC (S5b Table) to compare phage activity with different *E. coli* pathogenic strains. Overall, comparatively to ETEC, STEC and ETEC/STEC strains, FJ1, FN and SP1 were able to infect and propagate in a higher number of strains, respectively, 13.9%, 25.0% and 13.9% from which 100% (FJ1), 55.6% (FN) and 60% (SP1) had an EOP greater than 50% (LFW in 22.2%, 38.9% and 38.9%) of the APEC.

### Phage morphology and genome

TEM images showed that all phages are tailed (*Caudovirales* order) and belong to the *Myoviridae* family (Fig 2a). They have highly similar genomes ranging from 165 to 170 kb (269 to 280 coding sequences), sharing 87% overall nucleotide identity with *E. coli* phage T4, a prototype (NC_000866) of the subfamily *Tevenvirinae* (Fig 2b). Major phage proteins such as those related to DNA packing and structural proteins, DNA replication, recombination and modification proteins and cells lysis were identified in all genomes, however, >50% of the proteome has no assigned function. As expected, there was a high homology between the long tail fiber (LTF) of the three phages but less between them and the T4 phage (Fig 2c). A closer look at the distal part of the LTF indicated that such differences are mainly due to gp38 sequence (no homology found).

**Fig 2.**
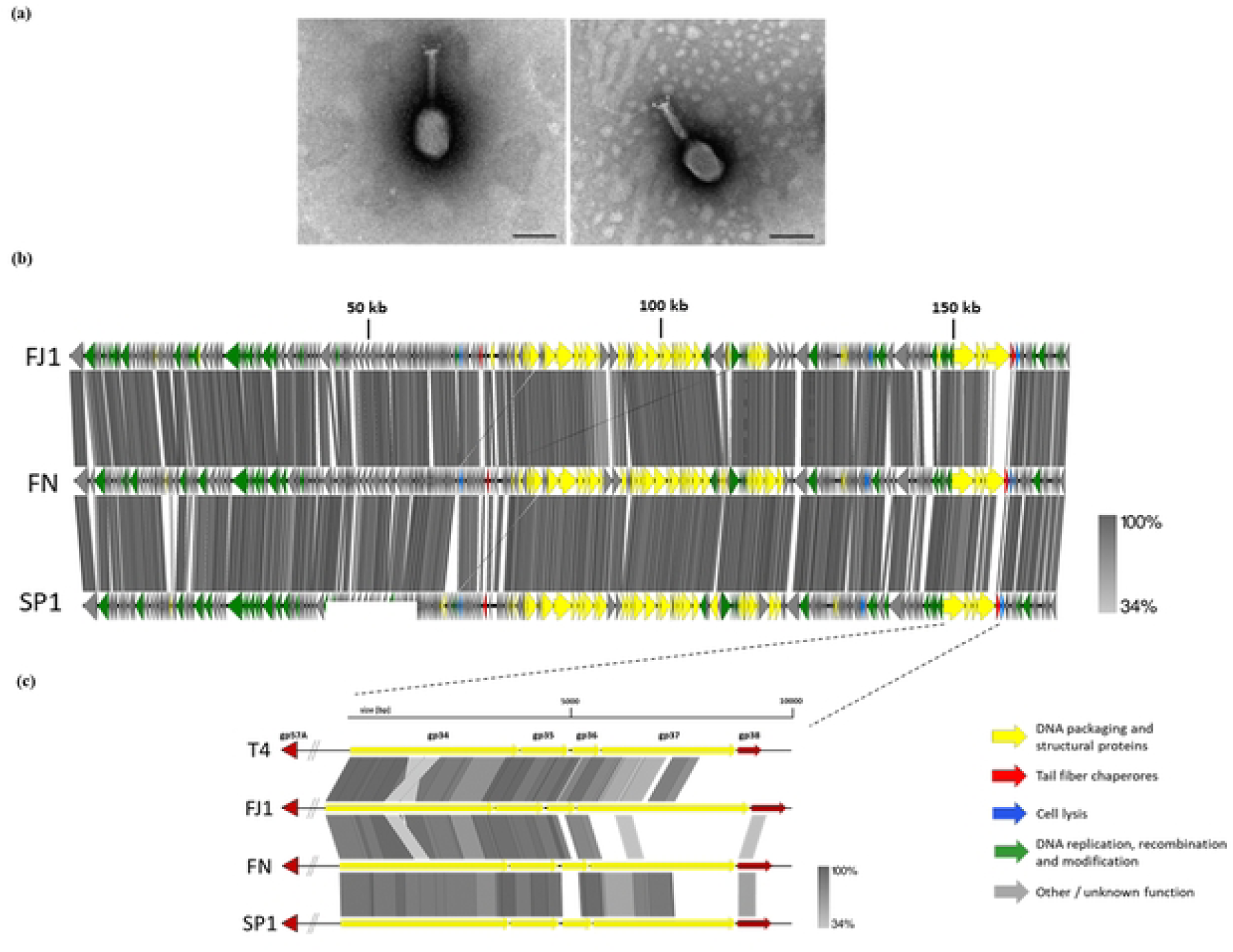
Microscopy observation and genomic comparison. **A)** Transmission electron micrographs of phage’s FJ1 (left) and FN (right). Black scale bar represents 100 nm. **B)** Comparison between the genomes of phage’s FJ1, FN and SP1. Coloured arrows indicate open reading frames according to the putative function. Similarity is indicated in grayscale. Image was created using the EasyFig program. **C)** Comparison between the putative coding sequences of the LTF of phage’s T4, FJ1, FN and SP1.

### Phage infection parameters

All three phages were evaluated in terms of one-step-growth curves (S1 Fig). Phages FJ1 and FN displayed the shortest latency periods of - 10 and 15 minutes, respectively - followed by SP1 that required 20 minutes to burst. Phages FN, FJ1 and SP1 produced 71, 96 and 150 phages per cell, respectively.

### Phage receptors

Preliminary assays aimed to infer about the nature of the phage receptors. Host cells were treated with periodate (to remove carbohydrates) or proteinase K (to remove outer membrane proteins) (Fig 3). When cells were treated with periodate, the adsorption rate significantly decreased for all phages (*p*-value< 0.001) - 9.8% ± 4.1% for FJ1, 45.8% ± 6.7 % for FN and 28.1% ± 9.1% for SP1 comparatively to untreated cells - 57.8% ± 7.3% for FJ1, 87.5% ± 3.5% for FN and 97.3% ± 1.8% for SP1. Adsorption of most phages was not affected when cells were treated with proteinase K, except for FN, with a reduction of 12%.

**Fig 3.**
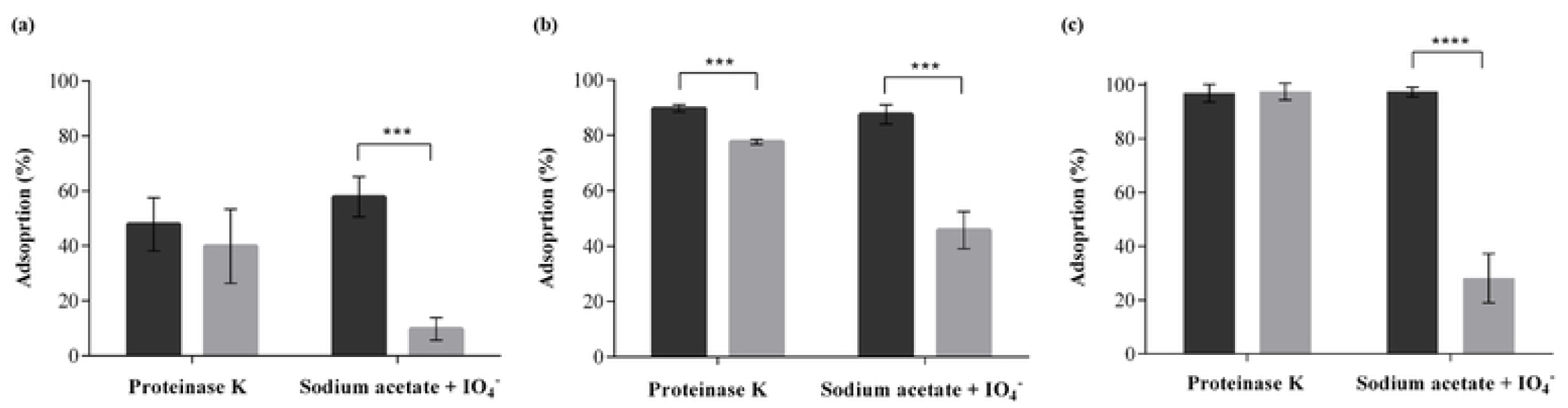
Adsorption assays of phages FJ1 (A), FN (B) and SP1 (C). Effect of proteinase K (0.2 mg.mL^−1^) and periodate (50 mM sodium acetate, pH 5.2, 100 mM IO_4_^−^) on phage host treated cells adsorption shown in residual PFU.mL^−1^ percentage. Controls were performed using distilled water instead of proteinase K or 50 mM sodium acetate, pH 5.2 only. Dark gray refers to control and light gray for treated cells. Errors bars represent standard deviation for an average of three repeated experiments. Significance was determined with t test when the treated and untreated groups were compared. *** *p*-value <0.001; **** *p*-value <0.001

Additional studies were conducted to detect the specific host receptor using a library of *E. coli* K-12 mutants. Results were in line with previous findings. Phage FN recognized proteins involved in the lipopolysaccharide (LPS) layer biosynthesis (RfaY, RfaG, RfaH and ADP-heptose--LPS heptosyltransferase 2 proteins) (either by moving sugar moieties and rearrange the structure of LPS or by enhancing the expression of operons involved in LPS synthesis) and binds to outer membrane proteins A (OmpA) as well. The specific receptor of phages FJ1 and SP1 was not possible to unveiled using the mutants tested.

### BIMs survival to serum antimicrobial activity

For this assay, 10 BIMs were confirmed and used for phages FJ1 and SP1, while only nine were obtained to FN (there was difficulty in obtaining any other, even after 72 h incubation). BIMs of the strain EC43 displayed different susceptibility towards the swine serum batches, depending on the originating phage (Fig 4). Overall, 90% of the FJ1 BIMs (1.1, 1.2, 1.3, 1.6, 1.8 and 1.9 (*p*-value <0.001), 1.4 (*p*-value=0.0021), 1.5 (*p*-value=0.0018) and 1.7 (*p*- value=0.0042)) and 100% of FN BIMs (2.1 (*p*-value=0.0060), 2.2 and 2.8 (*p*-value <0.001), 2.3 (*p*-value=0.0054), 2.4 (*p*-value=0.0027), 2.5 (*p*-value=0.0469), 2.6 (*p*-value=0.0024), 2.7 (*p*-value=0.0066), and 2.9 (*p*-value=0.0377)) were reduced, in average 1.2 ± 0.2 Log CFU.mL^−1^ comparatively to EC43 WT. Conversely, mutants generated after SP1 challenge did not show a higher susceptibility to serum killing activity when compared with the originating strain (*p-*value*>*0.05). The control tests performed with the inactivated serum did not influence the bacterial load concentration (data not shown), confirming that the reduction observed was only due to the bactericidal action of the serum.

**Fig 4.**
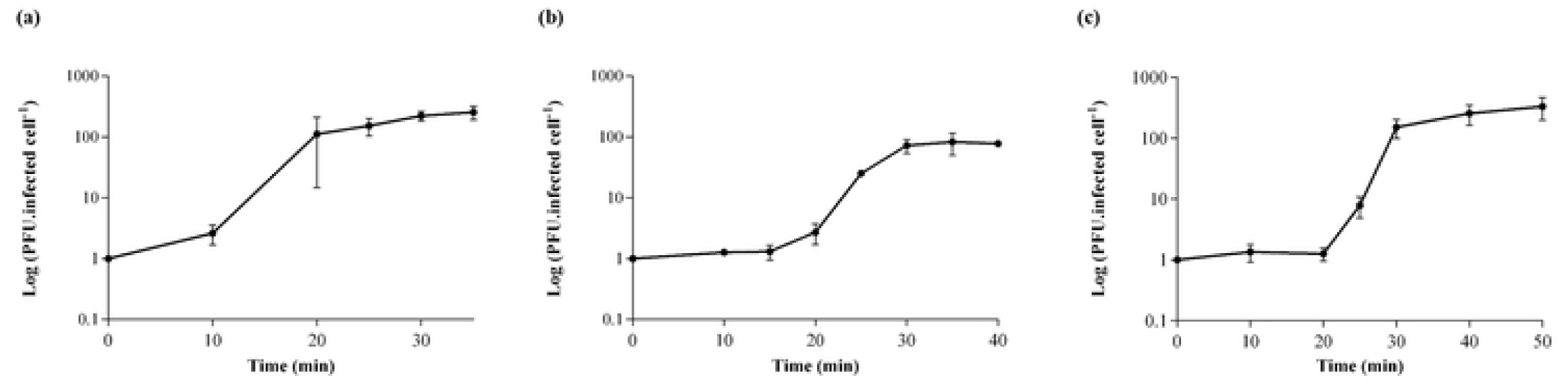
Serum complements effect against WT ETEC EC43 and respective BIMs. Mutants derived from interaction with phage’s FJ1 (**A** - 1.1, 1.2, 1.3, 1.4, 1.5, 1.6, 1.7, 1.8, 1.9, 1.10), FN (**B** - 2.1, 2.2, 2.3, 2.4, 2.5, 2.6, 2.7, 2.8 and 2.9) and SP1 (**C** - 3.1, 3.2, 3.3, 3.4, 3.5, 3.6, 3.7, 3.8, 3.9, 3.10). Porcine serum was challenged with EC43 WT and respective BIMs emerged from contact with each phage in a 3:1 ratio and incubated for 1 h at 37 °C. Results are shown in logarithm reduction of CFU.mL^−1^. Control was performed with inactivated serum (data not shown). Black refers to WT and dark gray for BIMs. Errors bars represent standard deviation for an average of three repeated experiments. Significance was determined with One-way ANOVA when the BIMs were compared with WT. * *p*-value <0.05; ** *p*- value <0.01; *** *p*-value <0.001; **** *p*-value<0.0001

### BIMs adhesion to intestinal porcine cell line

The mutants displayed different effect in the adhesion capacity to mammalian cells accordingly to the phage used to generated them (Fig 5). Most BIMs of FJ1 (80% - 1.1, 1.6 and 1.9 (*p*-value<0.001) and 1.4 (*p*-value=0.0102)) and FN (100% - 2.1 (*p*-value=0.0080), 2.2 and 2.9 (*p*-value<0.001), 2.3 (*p*-value=0.0022), 2.7 (*p*-value=0.0016)) demonstrated a reduced adhesion to culture cells, on average, of (5.4 ± 0.2) Log CFU.cm^−2^ and (5.2 ± 0.2) Log CFU.cm^−2^, correspondingly, comparing with the parental strain (6.3 ± 0.2) Log CFU.cm^− 2^. In opposition, SP1 generated BIMs did not show any difference regarding the adhesion capacity comparing with WT EC43 (*p*-value>0.05).

**Fig 5.**
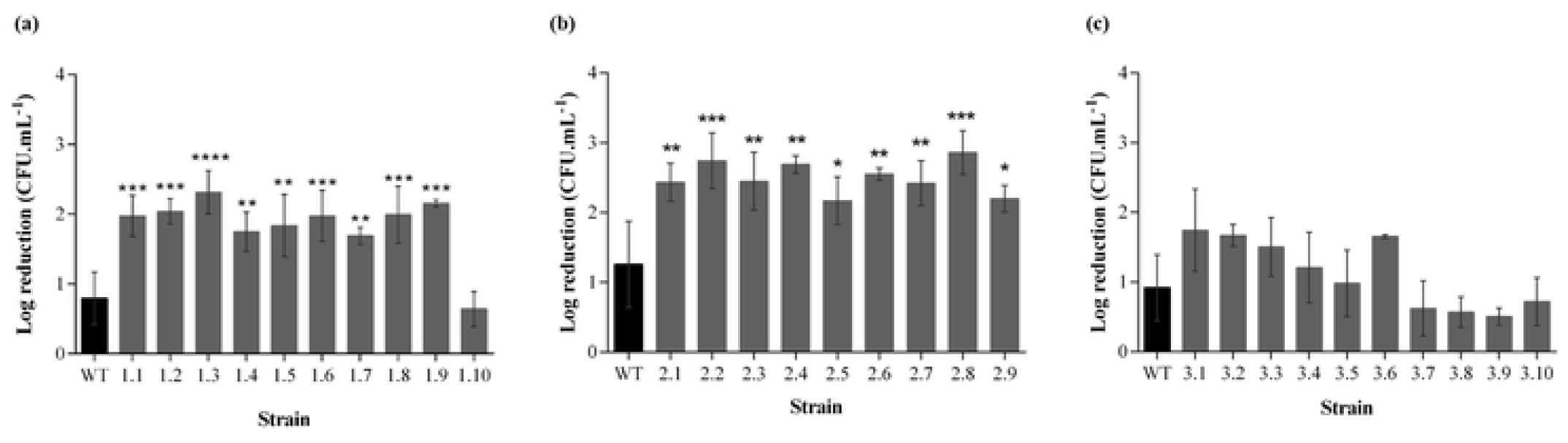
Adhesion capacity of WT ETEC EC43 and respective BIMs to mammalian cells. Mutants derived from interaction with phage’s FJ1 (**A** - 1.1, 1.4, 1.6, 1.9 and 1.10), FN (**B** - 2.1, 2.2, 2.3, 2.7 and 2.9) and SP1 (**C** - 3.1, 3.5, 3.6, 3.7 and 3.9). IPEC-1 cells were challenged with EC43 WT and five BIMs emerged from contact with each phage using a MOI of 100 and incubated for 2 h at 37 °C, 5% CO_2_. Results are shown in logarithm of CFU.cm^−2^. Black refers to WT and dark gray for BIMs. Errors bars represent standard deviation for at least three repeated experiments. Significance was determined with One-way ANOVA when the BIMs were compared with WT. * *p*-value <0.05; ** *p*-value <0.01; *** *p*- value <0.001; **** *p*-value<0.0001

## Discussion

The proliferation of pathogenic *E. coli* in the intestine of pigs during the nursing and PW period has a great cost for the swine industry. The overuse of antibiotics in recent decades has triggered serious problems associated with antibiotic resistance events, compromising the therapeutic solutions available to fight against multidrug resistant swine colibacillosis. This study intended to assess the potential of three phages that were here fully characterized FJ1, FN and SP1- to tackle DEC strains, using a panel of ETEC, STEC and ETEC/STEC isolated from pigs.

Phage taxonomic and genomic characterization indicated that all are T4-like phages of the *Tevenvirinae* subfamily within the *Myoviridae* family, which, as the other T-even phages, are known to infect a wide variety of Gram-negative hosts [24,25]. When host recognition and plaque formation assays were performed for each phage, all revealed an unexpected narrow host spectrum. As noted earlier [26], the number of infected phage- propagating strains was reduced compared to those which, despite being recognized and lysed, cannot propagate them: mostly low EOP scores (<50%) were recorded, and only in few strains: 1.0% for FJ1 and SP1 and 2.9% for FN, compared to the occurrence rates of LFW events, 4.8% for FJ1, 19.2% for FN and 9.6% for SP1. The wide variety of strains used not only the herein isolated but also the previously reported and characterized, isolated from Spanish farms [5] - contributed to the robustness and heterogenicity of this analysis. The panel comprised mostly fimbriae carriers’ strains (75%) belonging to 15 different serogroups (including the most prevalent within swine colibacillosis), with 94 ETEC and ETEC/STEC strains harbouring mostly toxin STb (76.6%), followed by STa (55.3%) and LT (42.6%) and 10 STEC. Moreover, the fact that 71.2% were *mcr*+, >40% carry beta-lactamase-encoding genes (including 18.3% extended-spectrum beta-lactamase producers) and 82.7% MDR strains reinforce the relevance of this work. It has been suggested that phages infecting but not being able to propagate in their hosts may be targeted by bacterial anti-phage defense systems, such as Restriction Modification (RM), or Abortive Infection (Abi) systems [26]. If the same report proposed that specifically T4-like phages might exhibit broad resistance to RM systems, it also indicated that they may be susceptible to some Abi systems, as typically their hosts encode genes able to sense phage specific proteins, triggering cell destruction and preventing subsequent infections. Analyzing now the low EOP observed, the comparison with APEC scores (higher EOP efficacy and wide host recognition, between 13.9% and 25%) seems to indicate (regardless of the phages isolation origin) that particularly ETEC may be strengthening its immunity against these viral predators. It can be speculated that this was possible, for example through host specialization in anti-phage defense systems such as CRISPR-CAS or Superinfection exclusion (SE) [27]. In an extensive *in silico* study, Wang et al. (2020) [28] demonstrated that ETEC can include 8.4 prophages/genome, a high average number, increasing the likelihood of occurring SE events.

T-even phages are known to infect hosts through an initial and reversible binding to primary receptors, - usually surface proteins like OmpC for T4, but when not available, sugar motifs in the LPS - with its LTF, followed but second and irreversible binding by its short tail fiber [29]. However, it is the first step that defines the host range. In phages such as T4 and S16, the LTF is encoded by gp34 to gp37 which form the tail proximal to distal segments [30,31] known to bind to both LPS moieties and OMP proteins. It is also known that T4 binds to hosts via gp37 but needs to be co-expressed with gp38 that functions as a chaperone [32], while S16 binds to hosts using gp37 bond with the gp38 which acts as an adhesin and mediates host specificity [31]. Given the fact that our *E. coli* phages share similar proximal end of the LTFs but have different distal segments (from the C-terminal gp37 to gp38), this can also explain their different and unexpected narrow host range. Regarding their host receptors, all the three phages are expected to be behave similarly to reported T-even phages. Accordingly, assays revealed that FN binds to proteins involved in LPS biosynthesis and OmpA. FJ1 and SP1 seems to recognize only one receptor (carbohydrates), nonetheless, we were not able to identify them using K-12 mutants. The repeated *in vitro* exposure of bacterial hosts to our phages led to the emergence of BIMs. This new phenotype brought fitness cost to some of the mutants, depending on the inducing phage. In fact, 90% of the BIMs from FJ1 and 100% from FN were more vulnerable to the pig complement system when compared to the originating strain, suggesting that changes at the level of the cell wall made the host more reactive to serum immunogenic proteins [33]. Conversely, pig serum has no increased bactericidal effect against BIMs from SP1, recognizing only LPS. Differences regarding serum sensitivity suggest that the site of the mutations at least at the LPS level will influence the reaction of the immune system. Mizoguchi and colleagues had already reported the formation of phage resistant strains associated with LPS alterations or OMP deficiency [34]. Particularly the surface protein OmpA (targeted by FN) and also carbohydrates (targeted by FJ1 and FN), especially those that are part of the LPS have been implicated in resistance to serum [35]. Furthermore, the bacterial structure re-conformation due to loss or alteration of the FJ1 and FN binding sites in ETEC EC43 influenced the adhesion capacity to porcine mammalian cells (in about 90%), even though the target of both phages is not related to their adhesin structures. This seems of relevant importance since the first step in the colonization of diarrhoeagenic pathotypes such as ETEC is their attachment to the host cells that promotes the transferring of enterotoxins more efficiently to the target cells.

## Conclusions

In summary, this study reflects the diversity of ETEC and STEC in Portuguese swine farms associated with high resistance towards several class of antibiotics, including last resort antimicrobials. Additionally, it demonstrated that the host range of three phages appears to be conditioned by the presence of a unique region in the phage’s LTFs. Overall, it suggests the importance of improving knowledge about ETEC and phage interaction, enhancing the importance of an extensive study of phage for a potential veterinary use.

## Supporting information

**S1 Table. Targets and primers associated with ETEC and STEC pathotypes**.

**S2 Table. Primers used for the detection and/or sequencing of TEM, SHV, CTX-M and MCR genes**.

**S3 Table. Targets and primers to determine phylogroups, clonotypes and sequence types by MLST**.

**S4 Table. Molecular characterization of the 24 *mcr*+ Portuguese strains**.

**S5 Table. Phages (FJ1, FN and SP1) lytic spectra and EOP against 104 ETEC, STEC and ETEC/STEC strains (a) and 36 APEC strains (b)**. The EOP was divided into four scores: 0 (no lysis), 1 (≤50%), 2 (>50% - 100%) and 3 (>100%). LFI stands for lysis from within and LFW for lysis from without. Light gray represents ETEC strains, gray cells are STEC strains and dark gray stands for ETEC/STEC isolates. NA: Not assigned, ONT: O non-typeable, HNM: H non motile.

**S1Fig.**
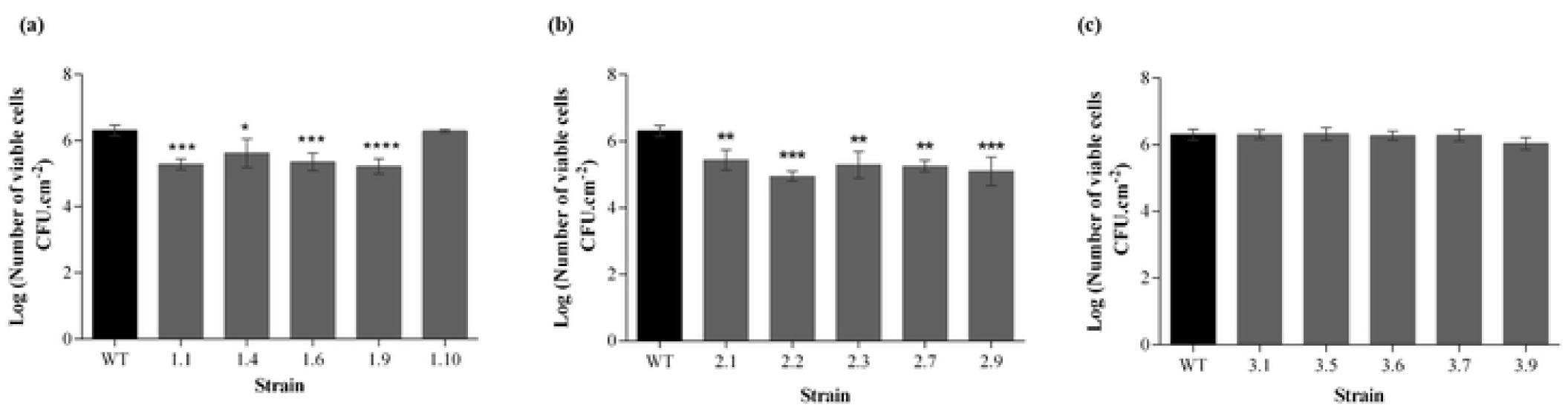
One-step growth curve of phages FJ1 (a), FN (b) and SP1 (c) on respective hosts cells. Results are shown in PFU per infected cell. Errors bars represent standard deviation for an average of three repeated experiments.

## References

1. Luppi A. Swine enteric colibacillosis: Diagnosis, therapy and antimicrobial resistance. Porc Heal Manag. 2017;3:1–18.

2. Mellor DJ, Stafford KJ. Animal welfare implications of neonatal mortality and morbidity in farm animals. Vet J. 2004;168(2):118–33.

3. Nagy B, Fekete PZ. Enterotoxigenic Escherichia coli in veterinary medicine. Int J Med Microbiol. 2005;295(6–7):443–54.

4. Luppi A, Gibellini M, Gin T, Vangroenweghe F, Vandenbroucke V, Bauerfeind R, et al. Prevalence of virulence factors in enterotoxigenic Escherichia coli isolated from pigs with post-weaning diarrhoea in Europe. Porc Heal Manag. 2016;2:1–6.

5. García-Meniño I, García V, Mora A, Díaz-Jiménez D, Flament-Simon SC, Alonso MP, et al. Swine enteric colibacillosis in Spain: Pathogenic potential of mcr-1 ST10 and ST131 E. Coli Isolates. Front Microbiol. 2018;9(11):1–15.

6. EMA. Categorisation of antibiotics in the European Union. European Medicines Agency (EMA/CVMP/CHMP/682198/2017). 2019. Available from: https://www.ema.europa.eu/en/documents/report/categorisation-antibiotics-european-union-answer-request-european-commission-updating-scientific_en.pdf Accessed 10 November 2021.

7. Salmond GPC, Fineran PC. A century of the phage: past, present and future. Nat Rev Microbiol. 2015;13(12):777–86.

8. Zhang J, Li Z, Cao Z, Wang L, Li X, Li S, et al. Bacteriophages as antimicrobial agents against major pathogens in swine: A review. J Anim Sci Biotechnol. 2015;6(1):1–7

9. Cha S Bin, Yoo AN, Lee WJ, Shin MK, Jung MH, Shin SW, et al. Effect of bacteriophage in enterotoxigenic Escherichia coli (ETEC) infected pigs. J Vet Med Sci. 2012;74(8):1037–9.

10. Jamalludeen N, Johnson RP, Shewen PE, Gyles CL. Evaluation of bacteriophages for prevention and treatment of diarrhea due to experimental enterotoxigenic Escherichia coli O149 infection of pigs. Vet Microbiol. 2009;136(1–2):135–41.

11. Lee CY, Kim SJ, Park BC, Han JH. Effects of dietary supplementation of bacteriophages against enterotoxigenic Escherichia coli (ETEC) K88 on clinical symptoms of post-weaning pigs challenged with the ETEC pathogen. J Anim Physiol Anim Nutr (Berl). 2017;101(1):88–95.

12. Mora A, Viso S, López C, Alonso MP, García-Garrote F, Dabhi G, et al. Poultry as reservoir for extraintestinal pathogenic Escherichia coli O45: K1: H7-B2-ST95 in humans. Vet Microbiol. 2013;167(3–4):506–12.

13. Clermont O, Christenson JK, Denamur E, Gordon DM. The Clermont Escherichia coli phylo-typing method revisited: Improvement of specificity and detection of new phylo-groups. Environ Microbiol Rep. 2013;5(1):58–65.

14. Wirth T, Falush D, Lan R, Colles F, Mensa P, Wieler LH, et al. Sex and virulence in Escherichia coli: An evolutionary perspective. Mol Microbiol. 2006;60(5):1136–51.

15. Weissman SJ, Johnson JR, Tchesnokova V, Billig M, Dykhuizen D, Riddell K, et al. High-resolution two-locus clonal typing of extraintestinal pathogenic Escherichia coli. Appl Environ Microbiol. 2012;78(5):1353–60.

16. Guinée, P. A. M., Jansen, W. H., and Wasdtröm TRS. Laboratory Diagnosis in Neonatal Calf and Pigs Diarrhoea: Current Topics in Veterinary and Animal Science. 1981;13 p. Netherlands: Springer.

17. Magiorakos A, Srinivasan A, Carey RB, Carmeli Y, Falagas ME, Giske CG, et al. Bacteria : an International Expert Proposal for Interim Standard Definitions for Acquired Resistance. 2011;18(3):268–81.

18. Ferreira A, Oliveira H, Silva D, Almeida C, Burgan J, Azeredo J, et al. Complete Genome Sequences of Eight Phages Infecting Enterotoxigenic Escherichia coli in Swine. Microbiol Resour Announc. 2020;9(36):8–11.

19. Melo LDR, Sillankorva S, Ackermann HW, Kropinski AM, Azeredo J, Cerca N. Isolation and characterization of a new Staphylococcus epidermidis broad-spectrum bacteriophage. J Gen Virol. 2014;95(Pt2):506–15.

20. Aziz RK, Bartels D, Best AA, DeJongh M, Disz T, Edwards RA, et al. The RAST Server: Rapid Annotations using Subsystems Technology. BMC Genomics. 2008;9(1):75.

21. Lowe TM, Eddy SR. TRNAscan-SE: A program for improved detection of transfer RNA genes in genomic sequence. Nucleic Acids Res. 1996;25(5):955–64.

22. Kiljunen S, Datta N, Dentovskaya S V., Anisimov AP, Knirel YA, Bengoechea JA, et al. Identification of the lipopolysaccharide core of Yersinia pestis and Yersinia pseudotuberculosis as the receptor for bacteriophage ΦA1122. J Bacteriol. 2011;193(18):4963–72.

23. Baba T, Ara T, Hasegawa M, Takai Y, Okumura Y, Baba M, et al. Construction of Escherichia coli K-12 in-frame, single-gene knockout mutants: The Keio collection. Mol Syst Biol. 2006;2:2006.0008.

24. Marti R, Zurfluh K, Hagens S, Pianezzi J, Klumpp J, Loessner MJ. Long tail fibres of the novel broad-host-range T-even bacteriophage S16 specifically recognize Salmonella OmpC. Mol Microbiol. 2013;87(4):818–34.

25. Xu J, Chen M, He L, Zhang S, Ding T, Yao H, et al. Isolation and characterization of a T4-like phage with a relatively wide host range within Escherichia coli. J Basic Microbiol. 2016;56(4):405–21.

26. Maffei E, Shaidullina A, Burkolter M, Heyer Y, Estermann F, Druelle V, et al. Systematic exploration of Escherichia coli phage–host interactions with the BASEL phage collection. PLOS Biology. 2021;19(11): e3001424.

27. Hampton HG, Watson BNJ, Fineran PC. The arms race between bacteria and their phage foes. Nature. 2020;577(7790):327–36.

28. Wang M, Zeng Z, Jiang F, Zheng Y, Shen H, Macedo N, et al. Role of enterotoxigenic Escherichia coli prophage in spreading antibiotic resistance in a porcine-derived environment. Environ Microbiol. 2020;22(12):4974–84.

29. Trojet SN, Caumont-sarcos A, Perrody E. Comeau AM, Krisch HM. The gp38 adhesins of the T4 Superfamily: a complex modular determinant of the phage’s host specificity. Genome Biol Evol. 2011;3:674–86.

30. King J, Laemmli UK. Polypeptides of the tail fibres of bacteriophage T4. J Mol Biol. 1971;62(3):465–77.

31. Dunne M, Denyes JM, Arndt H, Loessner MJ, Leiman PG, Klumpp J. Salmonella Phage S16 Tail Fiber Adhesin Features a Rare Polyglycine Rich Domain for Host Recognition. Structure. 2018;26(12):1573-1582.e4.

32. Hashemolhosseini S, Stierhof YD, Hindennach I, Henning U. Characterization of the helper proteins for the assembly of tail fibers of coliphages T4 and ?. J Bacteriol. 1996;178(21):6258–65.

33. Sumrall ET, Shen Y, Keller AP, Rismondo J, Pavlou M, Eugster MR, et al. Phage resistance at the cost of virulence: Listeria monocytogenes serovar 4b requires galactosylated teichoic acids for InlB-mediated invasion. PLoS Pathog. 2019;15(10):1–29.

34. Mizoguchi K, Morita M, Fischer CR, Yoichi M, Tanji Y, Unno H. Coevolution of bacteriophage PP01 and Escherichia coli O157:H7 in continuous culture. Appl Environ Microbiol. 2003;69(1):170–6.

35. Miajlovic H, Smith SG. Bacterial self-defence: How Escherichia coli evades serum killing. FEMS Microbiol Lett. 2014;354(1):1–9.

